# Mozambican genetic variation provides new insights into the Bantu expansion

**DOI:** 10.1101/697474

**Authors:** Armando Semo, Magdalena Gayà-Vidal, Cesar Fortes-Lima, Bérénice Alard, Sandra Oliveira, João Almeida, António Prista, Albertino Damasceno, Anne-Maria Fehn, Carina Schlebusch, Jorge Rocha

## Abstract

The Bantu expansion, which started in West Central Africa around 5,000 BP, constitutes a major migratory movement involving the joint spread of peoples and languages across sub-Saharan Africa. Despite the rich linguistic and archaeological evidence available, the genetic relationships between different Bantu-speaking populations and the migratory routes they followed during various phases of the expansion remain poorly understood. Here, we analyze the genetic profiles of southwestern and southeastern Bantu-speaking peoples located at the edges of the Bantu expansion by generating genome-wide data for 200 individuals from 12 Mozambican and 3 Angolan populations using ∼1.9 million autosomal single nucleotide polymorphisms. Incorporating a wide range of available genetic data, our analyses confirm previous results favoring a “late split” between West and East Bantu speakers, following a joint passage through the rainforest. In addition, we find that Bantu speakers from eastern Africa display genetic substructure, with Mozambican populations forming a gradient of relatedness along a North-South cline stretching from the coastal border between Kenya and Tanzania to South Africa. This gradient is further associated with a southward increase in genetic homogeneity, and involved minimum admixture with resident populations. Together, our results provide the first genetic evidence in support of a rapid North-South dispersal of Bantu peoples along the Indian Ocean Coast, as inferred from the distribution and antiquity of Early Iron Age assemblages associated with the Kwale archaeological tradition.

## Introduction

It is generally believed that the dispersal of Bantu languages over a vast geographical area of sub-Saharan Africa is the result of a migratory wave that started in the Nigeria-Cameroon borderlands around 4,000-5,000 BP (1–3). Although the earliest stages of the Bantu expansions were probably not associated with plant cultivation and domestication, Bantu speech communities added agriculture and iron metallurgy to their original subsistence strategies and subsequently replaced or assimilated most of the resident forager populations who lived across sub-Saharan Africa (4, 5). For this reason, the dispersal of Bantu-speaking peoples has often been considered a prime example of the role of food production in promoting demic migrations and language spread (6).

While genetic studies had a pivotal role in demonstrating that the Bantu expansions involved a movement of people (demic diffusion) rather than a mere spread of cultural traits (7–10), the majority of research on the specific routes and detailed dynamics of the spread of Bantu-speakers has been conducted in the fields of linguistics and archaeology.

Linguistic studies focusing on the reconstruction of the historical relationships between modern Bantu languages have led to some rather concrete proposals about links between individual languages and language areas, including the establishment of three widely accepted geographical subgroups: North-West Bantu, East Bantu and West Bantu (1, 11, 12). Among them, the East Bantu languages, which currently extend from Uganda to South Africa, have been shown to form a single monophyletic clade that is believed to be a relatively late offshoot of West Bantu (13–15). Assuming that the phylogenetic trees inferred from the comparison of lexical data can be used to trace the migratory routes of ancestral Bantu-speaking communities, the linguistic pattern favors a dispersal scenario whereby populations from the Nigeria-Cameroon homeland first migrated to the south of the rainforest and later diversified into several branches before occupying eastern and southern Africa (14, 15).

According to archaeological evidence, the earliest Bantu speakers in East Africa appeared around 2,600 BP in the Great Lakes region, associated with pottery belonging to the so-called Urewe tradition, also characterized by a distinctive iron smelting technology and farming (1, 16). However, the link between Urewe and pottery traditions further west is unclear, and the historical events leading to its introduction to the interlacustrine area are still poorly understood (17). Some interpretations of the archaeological data have proposed that, in contrast with the “late split” between East and West Bantu suggested by linguistic evidence, East Bantu peoples introduced the Urewe tradition into the Great Lakes by migrating out of the proto-Bantu heartland along the northern fringes of the rainforest after an early separation from Bantu speakers occupying the western half of Africa (18, 19). This model, however, is not supported by recent genetic studies showing that Bantu-speaking populations from eastern and southern Africa are more closely related to West Bantu speakers that migrated to the south of the rainforest than they are to West Bantu speakers that remained in the north (20–22).

In spite of their uncertain origins, the Urewe assemblages display pottery styles similar to the younger Kwale and Matola traditions that are distributed along coastal areas ranging from southern Kenya across Mozambique to KwaZulu-Natal (1, 16, 17, 23). This archaeological continuity has been interpreted as the earliest material evidence for an extremely rapid dispersion of East Bantu speakers from the Great Lakes, starting around the second century AD and reaching South Africa in less than two centuries (1, 16, 23). Such a migration remains, however, to be documented by genetic data, due to insufficient sampling of the areas lying between eastern and southern Africa that roughly correspond to present-day Mozambique.

In this study, we fill this important gap by investigating the population history of Mozambique using ∼1.9 million quality-filtered single nucleotide polymorphisms (SNPs) that were genotyped in 161 individuals from 12 populations representing all major Mozambican languages, and in 39 individuals from 3 contextual populations from Angola (Fig. 1 and Table S1). By making use of a maximally wide range of available genetic and linguistic data, we show that East Bantu-speaking populations display genetic substructure, and detect a strong signal for the dispersal of East Bantu peoples along a North-South cline, which possibly started in the coastal border between Kenya and Tanzania and involved minimum admixture with local foragers until the Bantu-speakers reached South Africa. Together, our results provide a strong support for reconstructions of the eastern Bantu migrations based on the distribution of Kwale archaeological sites.

**Fig. 1.**
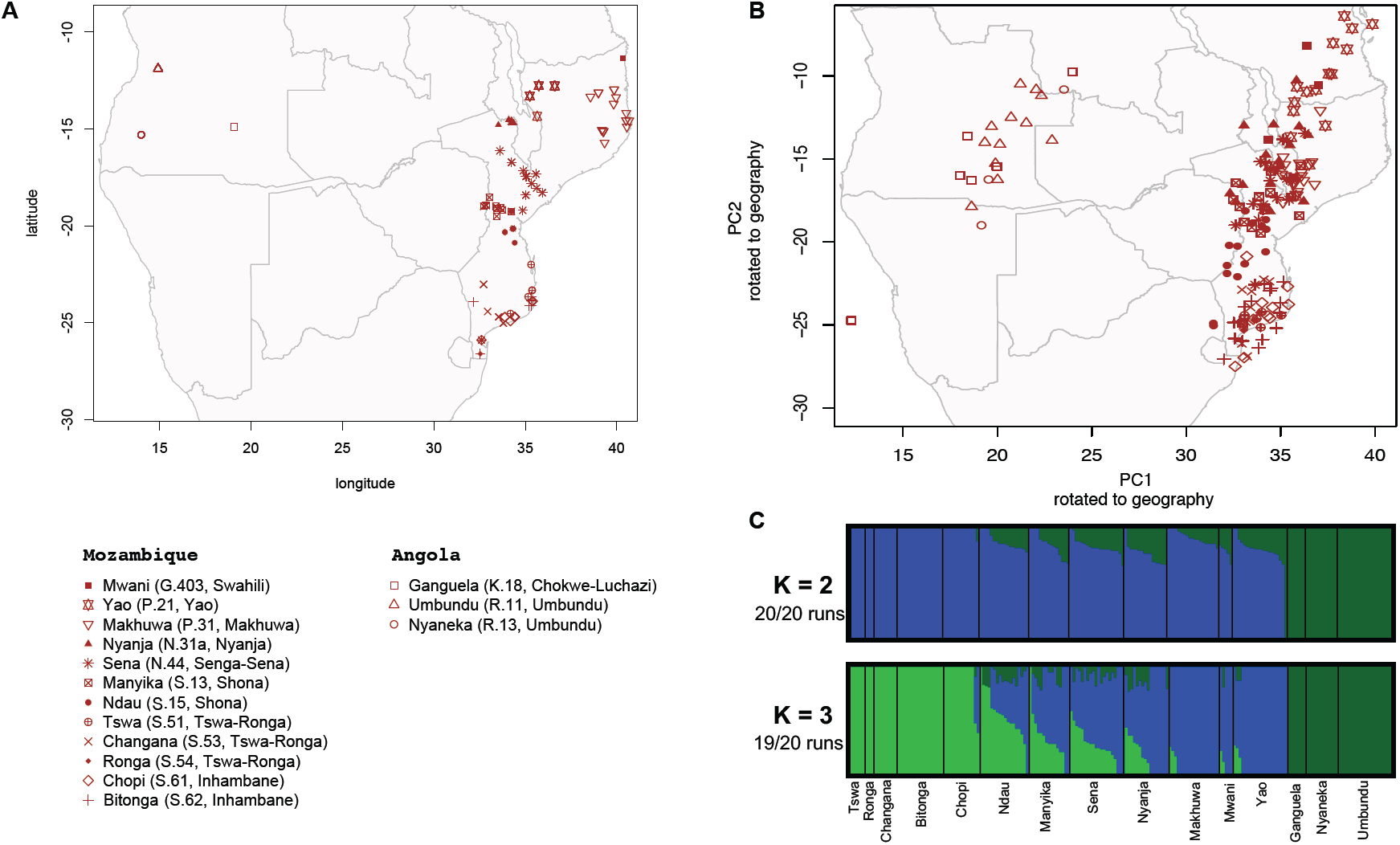
Genetic structure in Angolan and Mozambican populations. (A) Geographic locations of sampled individuals. The geographic subgroups of Bantu languages (“Guthrie zones”) following Maho (26) are given in parentheses in the legend. (B) Principal components 1 and 2 of Angolan and Mozambican individuals rotated to fit geography (Procrustes correlation: 0.89; P<0.001). (C) Population structure estimated with ADMIXTURE assuming 2 and 3 clusters (K). Vertical lines represent the estimated proportion of each individual’s genotypes that are derived from the assumed genetic clusters (note that the order of individuals in K=2 is not the same as K=3). The lowest cross-validation error was associated with K=2.

## RESULTS AND DISCUSSION

### Genetic variation in Mozambique

To assess the genetic relationships between Angolan and Mozambican individuals, we performed principal component analysis (PCA) (24) and unsupervised clustering analysis using ADMIXTURE (25) (Fig. 1).

The PCA patterns are closely related to geography, with the first PC (PC1) separating Mozambican and Angolan individuals, and the second PC (PC2) revealing a noticeable heterogeneity among samples from Mozambique (Figs. 1*B* and S1; Procrustes correlation: 0.89; P<0.001). The ADMIXTURE analysis confirmed the substantial differentiation between populations from Angola and Mozambique (at K=2), and the genetic substructure among Mozambican populations (at K=3) (Fig. 1*C*).

Within Mozambique, the association between genetic patterns and geography is further highlighted by a strong correlation between average PC2 scores and latitude (r=-0.97, P<10^-6^), showing that genetic variation is structured along a North-South cline corresponding to the orientation of the country’s major axis (Fig. 2*A*). The highest genetic divergence was found between Yao and Mwani speakers in the north, and Tswa-Ronga (Tswa, Changana, Ronga) and Inhambane (Bitonga and Chopi) speakers in the south, while Makhuwa, Sena, Nyanja and Shona (Manyika and Ndau) speakers occupy intermediate genetic and geographic positions (Figs. 1*B* and 2*A*). Qualitatively, this trend is consistent with the geographic distribution of subclusters of Mozambican languages in the Bantu phylogeny proposed by Grollemund et al. (14) (cf. their Fig. S1). Our own lexicostatistical analyses reveal significant correlations between genetic and language pairwise distances (Mantel test: r=0.68; P=2.9 × 10 ^-5^), as well as between language and latitude (Mantel test: r=0.79; P=6.3 × 10 ^-5^) (Figs. S2; Tables S2-S4).

**Fig. 2.**
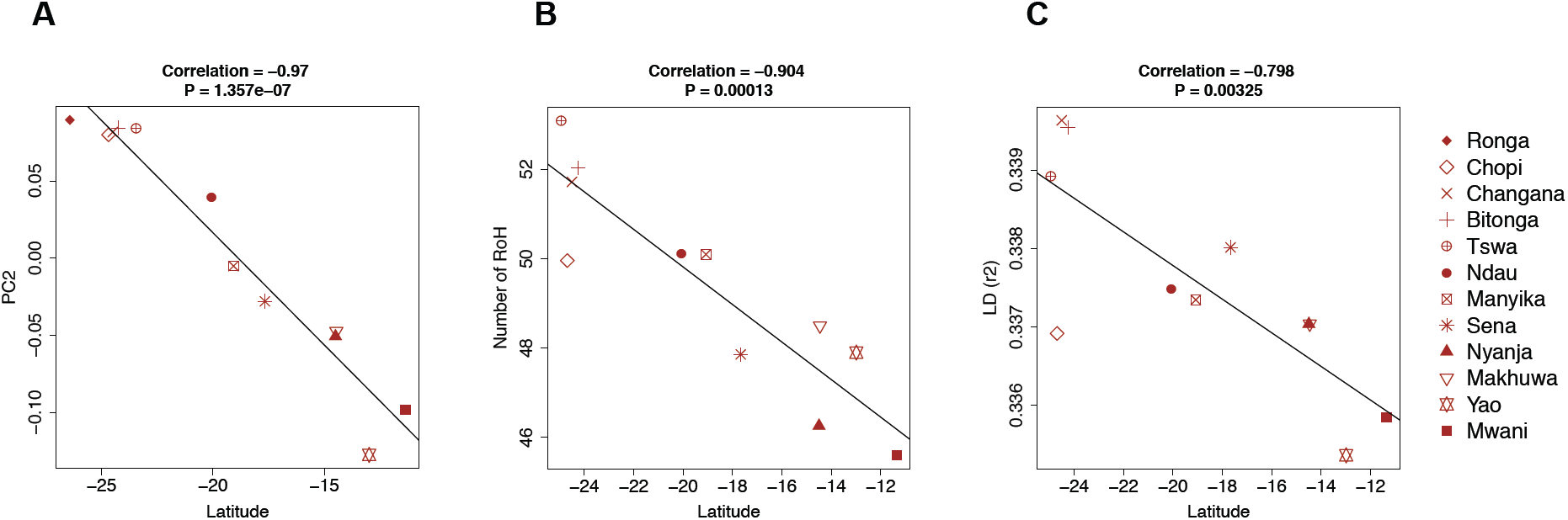
Genetic variation and geography in Mozambique. The plots show the correlations between latitude and (A) average PC2 scores (Fig. S1*B*) (B) average number of RoHs, and (C) average LD (r^2^). In B and C, Tswa and Ronga were lumped and are identified by the Tswa symbol (see SI Appendix).

A stepwise reduction in levels of genetic diversity with increasing geographic distance from a reference location is generally considered to be the typical outcome of a demic migration involving serial bottlenecks (27). In the global context of the Bantu expansion, a significant decrease of genetic diversity with distance to the Bantu homeland was previously reported for mitochondrial DNA and the Y-chromosome, but not for autosomes (9). Moreover, to our knowledge, there have been no reports for such patterns at more local scales. In order to evaluate the relationship between genetic diversity and geography, we studied the distribution of haplotype heterozygosity (HH), numbers and total lengths of runs of homozygosity (RoHs) and linkage disequilibrium (LD), as measured by the squared correlation of allele frequencies (*r*^*2*^), across all sampled Mozambican populations (*SI Appendix*).

We found that RoHs and LD were significantly correlated with latitude, with northern populations displaying higher genetic diversity than southern populations (Figs. 2*B* and *C*; Figs. S3-S5). We also observed a decrease of HH with absolute latitude that did not reach significance (Fig. S3*A*; r=0.51, P=0.104). However, HH was still significantly correlated with LD (Fig. S4*C*). Together, these results suggest that East Bantu-speaking peoples entered Mozambique from the North and underwent sequential reductions in effective population size, leading to increased genetic homogeneity and differentiation as they moved southwards.

To further assess the relationship between population structure and geography in Mozambique, we used the Estimated Effective Migration Surfaces (EEMS) method, which identifies local zones with increased or decreased migration rates, relative to the global migration across the whole country (28) (Fig. 3*A*). We detected two zones of low migration between northern and central Mozambique (Fig. 3*A*): one associated with Yao speakers, located in the northwestern highlands of the Nyasa Province between lake Nyasa/ Malawi and the Lugenda River (Figs. 3*B* and *C*); the other, located in the Northeast, to the north of the Ligonha River, around Makhuwa-speaking areas (Figs. 3*A* and *B*). An additional low-migration zone was found around the Save River, between southern and south-central Mozambique (Figs. 3*A* and *B*). Interestingly, the EEMS analysis also shows that the Zambezi River in central Mozambique is not an obstacle but rather a corridor for migration (Figs. 3*A* and *B*). Overall, the geographic patterns revealed by the EEMS method are consistent with the PC cline in showing that the highest genetic differentiation between the northernmost and southernmost populations is reinforced by intervening low migration zones, while the relative genetic proximity between central Mozambican groups was enhanced by increased migration around the Zambezi Basin (Figs. 3*A* and *B*).

**Fig. 3.**
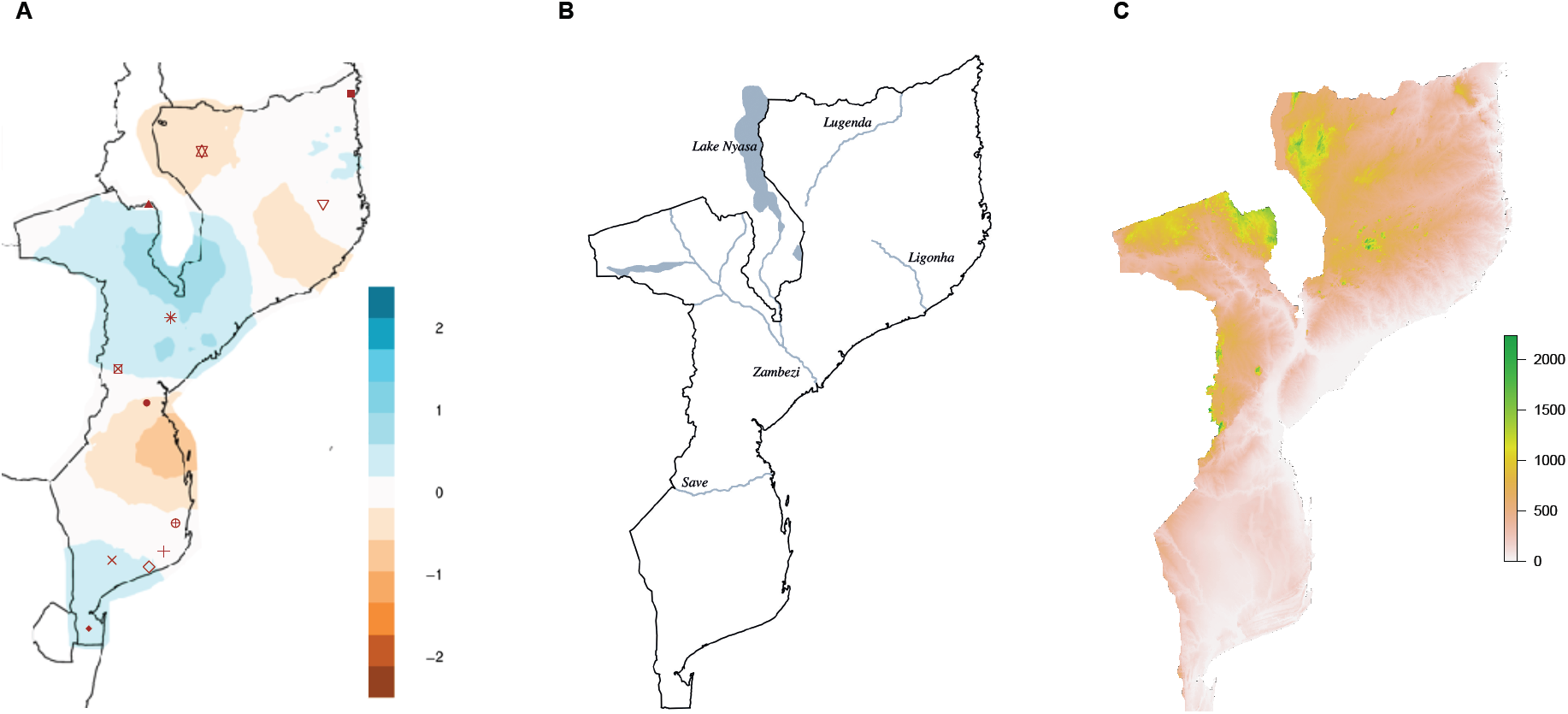
Estimated Effective Migration Surface (EEMS) analysis. See Fig. 1 for legend of population symbols. (A) EEMS estimated with 12 Mozambican populations. (B-C) Major rivers (B) and mountains (C) associated with barriers and corridors of migration. The effective migration rates are presented in a log10 scale: white indicates the mean expected rate in the dataset; blue and brown indicate migration rates that are X-fold higher or lower than average, respectively. The orographic map (C) was generated with the raster package (29). Altitude is given in meters.

### Genetic relationships with other African populations

To place the genetic variation of Mozambican and Angolan samples into the wider context of the Bantu expansion, we combined our dataset with available genome-wide comparative data from other African populations (Fig. 4*A* and Table S5).

**Fig. 4.**
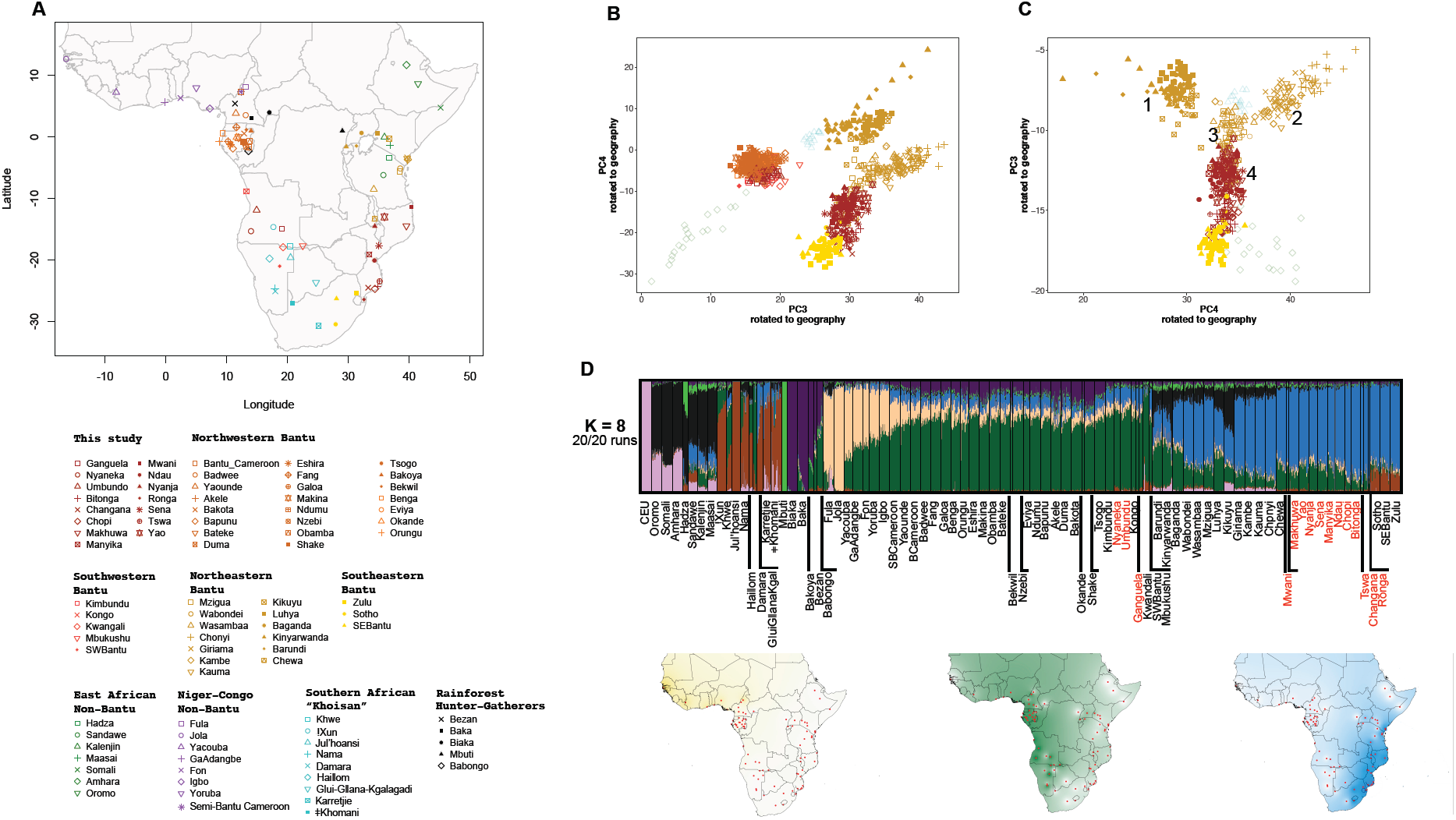
Genetic structure in African populations. (A) Geographic locations of sampled populations. (B-C) PC plots rotated to geography using Procrustes analysis. (B) All Bantu-speaking populations (Procrustes correlation: 0.76; P< 0.001). (D) Only East Bantu-speaking populations (Procrustes correlation: 0.44; P<0.001). The numbers in (C) refer to groups of populations that are discussed in the text. Additional PCA and ADMIXTURE plots are shown in Figs. S6 and S8. (D) Population structure estimated with ADMIXTURE assuming 8 clusters (K=8), with Mozambican and Angolan groups from this study labeled in red. Vertical lines represent the estimated proportions of each individual’s genotypes that are derived from the assumed genetic clusters. The maps, obtained by interpolation, display the mean proportions of major ADMIXTURE components from Niger-Congo-speaking populations. The colors in the maps match the colors in the ADMIXTURE plot.

Genetic clustering analysis shows that three partially overlapping components can be roughly associated with major geographic areas and linguistic subdivisions of the Niger-Congo phylum, of which the Bantu languages form part (Figs. 4*D* and S6): non-Bantu Niger-Congo in West Africa, to the north of the rainforest (beige); West Bantu, including Angolans, along the Atlantic coast (green); and East Bantu, including Mozambicans, in East Africa and along the Indian Ocean Coast (blue). A pairwise F_*st*_ analysis of all Niger-Congo speaking populations further shows that the highest levels of genetic differentiation are found between non-Bantu Niger-Congo groups and East Bantu-speaking peoples (Fig. S7).

Other major genetic components revealed by clustering analysis are associated with Kx’a, Tuu and Khoe-Kwadi-speaking peoples from southern Africa, also known as Khoisan (brown), Rainforest Hunter-Gatherers (RHG) (violet and light green), non-Bantu Eastern Africans (black) and Europeans (pink). As found in previous works (7, 20, 30), several Bantu-speaking populations have varying proportions of these genetic components, which were likely acquired through admixture with local residents: 11% (range: 4-21%) of RHG-related component in West Bantu speakers; 16% (range: 9-38%) of non-Bantu eastern African-related component in East Bantu speakers from Kenya and Tanzania; and 17% (range: 16-18%) of Khoisan-related component in southeastern Bantu speakers from South Africa.

To mitigate the effect of admixture with resident populations, we carried out a PC analysis of all Bantu-speaking groups, together with one representative group of non-Bantu Eastern Africans (Amhara) and one representative group of southern African Khoisan (Ju|’hoansi), which are the two most important sources for external admixture with Bantu-speaking populations from the East and South, respectively. As expected, the first two principal axes are driven by genetic differentiation between the Amhara (PC1) and the Ju|’hoansi (PC2), relative to Bantu-speaking groups (Fig. S8*A*). Moreover, some Bantu peoples from eastern (e.g., Kikuyu and Luhya) and southern Africa (e.g., Sotho and Zulu) stand out from a tight cluster encompassing all Bantu speakers by extending toward the Amhara and Ju|’hoansi, respectively, indicating admixture of local components into the genomes of Bantu-speaking populations. When the first two principal components are removed, a close link between the internal differentiation of Bantu-speaking groups and geography becomes apparent in a PC3 vs. PC4 plot (Figs. 4*B* and S8*E*; Procrustes correlation: 0.76; P< 0.001). In this plot, PC3 represents an east-west axis displaying a noticeable gap between West and East Bantu speakers, and PC4 highlights the differentiation of Mozambican and South African groups from eastern African populations located to their north. As shown by the PC3 vs. PC4 plot displayed in Fig. 4*C*, the heterogeneity of East Bantu populations is further emphasized when West Bantu speakers are removed (Procrustes correlation: 0.44; P< 0.001; Fig. S8*F*). While PC4 is correlated with longitude (r=-0.73; P<10^-4^), PC3 is highly correlated with latitude (r=0.95; P<10^-13^), showing that the gradient of genetic differentiation previously observed within Mozambique extends from eastern to southern Africa (Figs. 2*A* and S9). Heuristically, the genetic differentiation among East Bantu speakers can be described by defining four groups that are broadly associated with different geographic regions in eastern and southeastern Africa, and partially correspond to various linguistic zones of Guthrie’s Bantu classification (11, 26) (Figs. 4*C*, S9 and S10): 1) the first group includes peoples from the western fringe of eastern Africa (Kikuyu, Luhya, Baganda, Barundi and Kinyarwanda), who live around Lake Nyanza/Victoria and mostly speak languages belonging to Bantu zone J (Lakes Bantu) (31); 2) the second group includes populations from coastal Kenya (Chonyi, Giriama, Kambe and Kauma), who belong to the Mijikenda ethnic group and speak languages from zone E; 3) the third group is genetically intermediate between groups 1 and 2, and includes the Mzigua, Wabondei and Wasambaa from Tanzania, who speak languages from zone G; 4) the fourth group, formed by Mozambicans and South Africans, is an heterogeneous set of linguistically related populations covering zones N, P and S, who bridge the area between eastern and southern Africa and are genetically closer to groups from Tanzania than to other East Africans.

These findings have important implications for integrating archaeological, linguistic and genetic data in the reconstruction of the Bantu migrations in the easternmost regions of Africa. Although many crucial areas still need to be included in genome-wide analyses, the available data suggests that the occupation of eastern Africa by Bantu-speaking populations was associated with genetic structuring in the relatively small area between the Great Lakes and the Indian Ocean Coast, with Tanzanian Bantu-speaking populations providing the likely starting point for the chain of genetic differentiation events linked to the migration of Bantu-speaking peoples from eastern to southern Africa. This scenario is in close agreement with the migratory path inferred from the continuity between Early Iron Age (EIA) archaeological sites from the Kwale ceramic tradition, which extend from coastal Kenya and Tanzania to South Africa across a Mozambican corridor (1, 16, 23).

To further investigate the origins of the migratory streams linking different Bantu-speaking groups and to better characterize the admixture dynamics between Bantu speakers and resident populations, we applied the haplotype-based approaches implemented in CHROMOPAINTER and GLOBETROTTER (32, 33). We found that the haplotype copy profiles of Angolans differ significantly from Mozambicans+South Africans (Figs. 5*A* and *B*): while the former derive most of their haplotypes from West Bantu-speaking populations located to their North, the latter trace most of their ancestry to Bantu-speaking groups from East Africa, in close agreement with the PCA results (Fig. 4). More specifically, we found that the best donor population proxy (Mzigua) for Bantu speakers from Mozambique and South Africa is located in Tanzania (range: 72-93%), whilst Angolans derive most of their ancestry from Bantu-speaking groups in Gabon and Cameroon (range: 77-83%) (Fig 5*C*; Table S6).

**Fig. 5.**
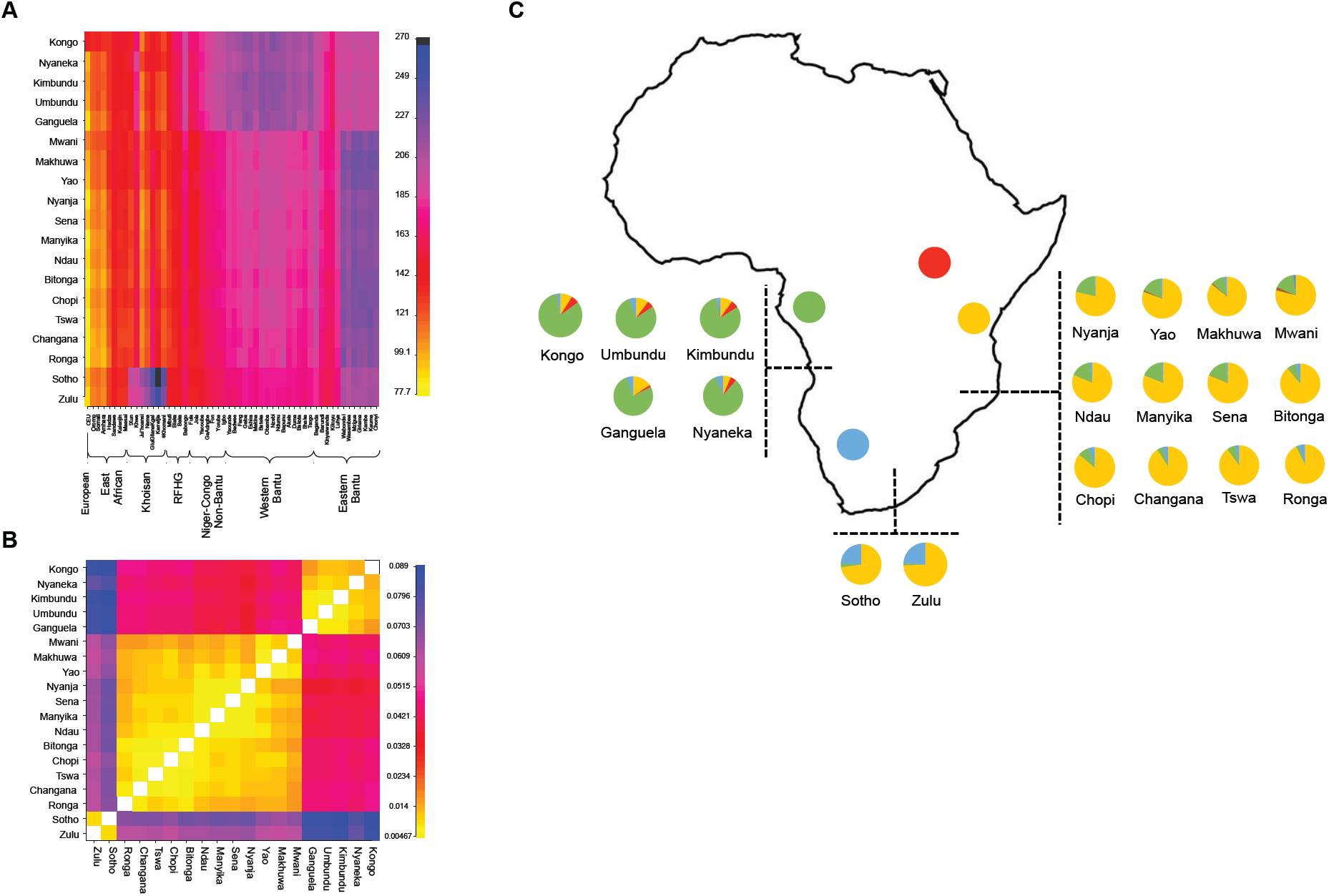
Inferred ancestry of Bantu-speaking groups from Angola, Mozambique and South Africa. (A) CHROMOPAINTER coancestry matrix based on the number of haplotype segments (chunk counts) shared between representative donor groups (columns) and recipient populations (rows) from Angola, Mozambique and South Africa. The copy profile of each recipient group is an average of the copy profiles of all individuals belonging to that group. (B) Matrix of pairwise TVDxy values based on the ancestry profiles of Angolan, Mozambican and South African groups. The scales of chunk counts and TVDxy values are shown to the right of the matrices in (A) and (B), respectively. (C) Ancestry profiles of Angolan, Mozambican and South African populations (pie charts) as inferred by the MIXTURE MODEL implemented in GLOBETROTTER. The colored circles indicate the most important contributing regions where best source populations were found: West Bantu-speaking groups (green); Tanzanian East Bantu-speaking groups (yellow); Great Lakes Bantu-speaking groups (red); and Khoisan groups (blue).

Estimated Khoisan ancestry in the South African Sotho (24%) and Zulu (24%) is much higher than in their close Mozambican neighbors Ronga (5%) and Changana (4%), or in any other Mozambican group (range: 1-5%) (Figs. 5*B* and *C*; Table S6). This pattern suggests that Bantu speakers scarcely admixed with local foragers, in agreement with recent findings about Bantu speakers from Malawi, who displayed no Khoisan ancestry, despite the confirmed presence of a Khoisan-related genetic component in ancient samples from the region (34). It therefore seems that the processes governing earlier admixture events between Bantu-speakers and local hunter-gather groups in modern-day Mozambique and Malawi were very different from what has been reported for South Africa and Botswana (7, 30, 35). As previously suggested on the basis of genetic variation in uniparental markers and archaeological modeling, the differences in admixture dynamics leading to increased Bantu/Khoisan admixture beyond the southern border of Mozambique could have been caused by a slowdown of the Bantu expansion due to adverse ecoclimatic conditions (36).

A recent genome-wide study found that the best-matching source population for South African Bantu speakers is located in Angola (Kimbundu) rather than in East Africa (as represented by the Bakiga and Luhya from around the Great Lakes) (20). Here, we used a stepwise approach to rank the best proxies for the ancestry of two South African Bantu-speaking groups (Sotho+Zulu) among all populations contained in our dataset (Fig. 6; SI Appendix; Table S6). We found that the Changana and Ronga from Mozambique, and a southern Khoisan descendent group (the Karretjie People of South Africa) are the best proxies for the ancestry of the South African Bantu speakers (Fig. 6*A*). When Mozambican populations are removed from the list of sources, the next best non-Khoisan proxies are the Mzigua from Tanzania (Fig. 6*B*). The contribution of Angola only becomes increasingly more relevant when Tanzanian (Fig. 6*C*), Kenyan (Fig. 6*D*) and Great Lakes (Fig. 6*E*) populations are successively removed from the list of donors. Nevertheless, the fact that Angola still represents a better proxy for the ancestry of southeastern Bantu speakers than populations closer to the Bantu homeland provides additional evidence in favor of a “late-split” between southwestern and southeastern Bantu-speaking groups after a single passage through the rainforest, as suggested in previous studies (20, 21).

**Fig. 6.**
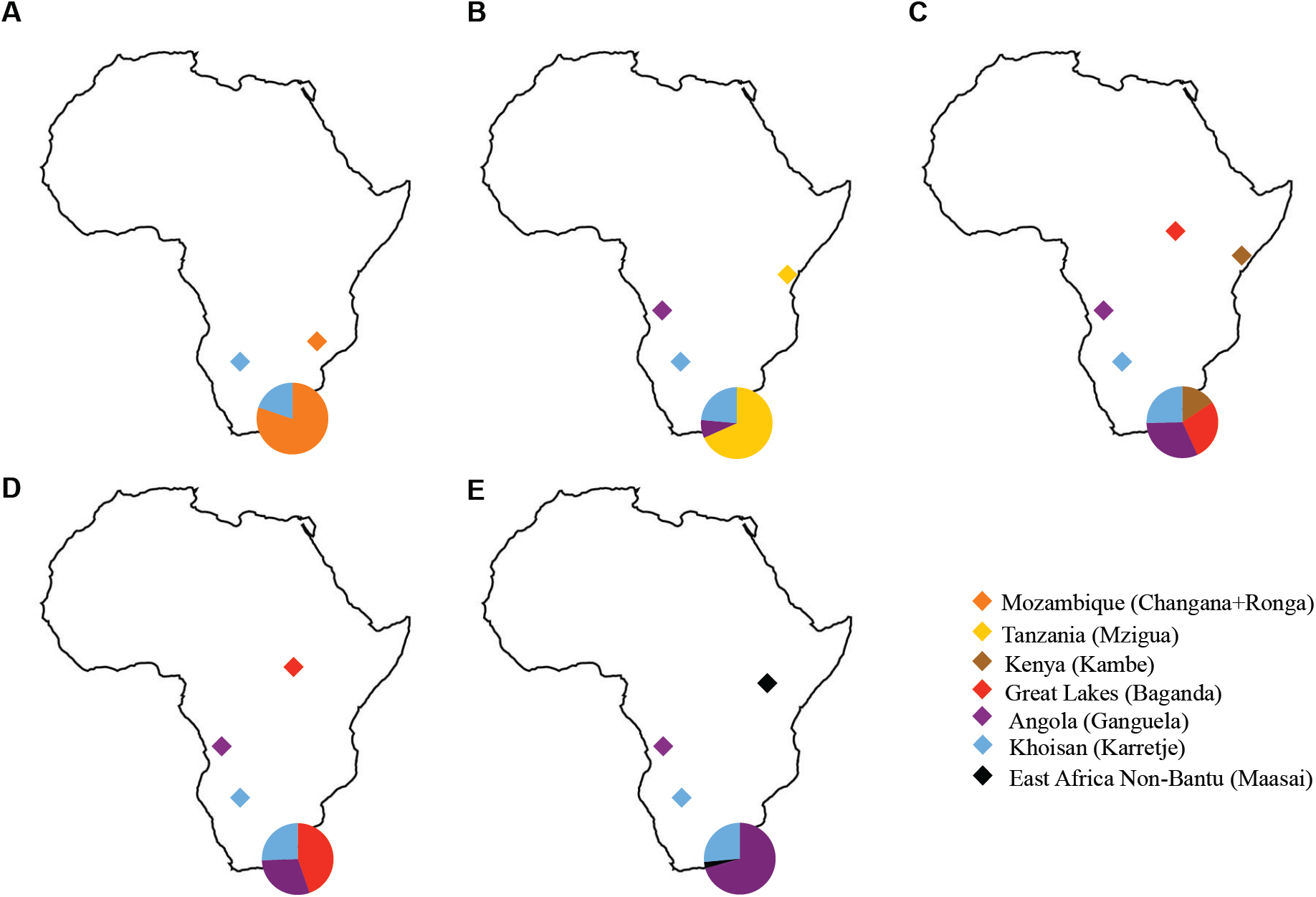
Inferred average ancestry of Bantu-speaking groups from South Africa. The most important contributing regions and best source populations are provided in the legend. (A) 71 source populations from Sub-Saharan Africa. (B) As in (A), but removing Mozambique from the list of sources. (C) As in (B), but removing Tanzanian Bantu speakers from the list of sources. (D) As in (C), but removing Bantu speakers from coastal Kenya from the list of sources. (E) As in (D) but removing Bantu speakers from the Great Lakes from the list of sources. Full lists of source populations are provided in Table S6.

In a further step, we identified and dated signals of admixture in the history of the studied populations using GLOBETROTTER. We found no evidence for admixture between any two Mozambican populations (not shown), suggesting that the intermediate position of central Mozambique in the North-South gradient of genetic relatedness (Figs. 1*B* and 2*A*) is not the result of admixture between populations from northern and southern Mozambique but rather a cline of stepwise genetic differentiation. At the same time, we found that the Khoisan ancestry detected in South Africans (Sotho and Zulu) and at low frequencies in southern Mozambican populations (Ronga, Changana, Tswa, Bitonga and Chopi) (Fig. 5*C*) resulted from admixture events occurring around 1165 BP (range: 756-1851 BP), involving the Karretjie people from South Africa as best matching Khoisan source and the Tanzanian Mzigua as best-matching Bantu-speaking population (P-values for evidence of admixture <0.05; Table S7). This date is remarkably consistent with the first Iron Age arrivals to southern Mozambique associated with the Matola pottery, which stylistically resembles the Kwale ceramics from Tanzania and has been dated to the early and mid-first millennium AD (23).

We also found evidence (P<0.05) for non-Bantu ancestry in Bantu speakers from the Great Lakes, coastal Kenya and Tanzania resulting from admixture with Afro-Asiatic (Amhara and Oromo) and Nilotic (Kalenjin and Maasai) speakers (Table S7). The average estimated antiquity of these admixture events dates to ∼760 BP (570-1047 BP) and is in close agreement with Bantu/non-Bantu eastern African admixture dates inferred by Skoglund et al. (34). These estimates postdate the Bantu/Khoisan admixture inferred for Mozambique and South Africa, suggesting that the bulk of admixture between Bantu and non-Bantu speakers in East Africa occurred only after Bantu speakers had already begun their migration towards the South. This is also supported by the low eastern African ancestry detected in Bantu speakers from Mozambique and South Africa.

## Conclusion

Using a country-wide sample of 12 Mozambican populations, we were able to fill an important gap in the understanding of the expansion of Bantu speakers from the Great Lakes region to the eastern half of southern Africa. Our results suggest that, in spite of the present-day homogeneity of East Bantu languages, the arrival of Bantu-speaking groups in eastern Africa was associated with a period of genetic differentiation in the area between the Great Lakes and the Indian Ocean Coast, followed by a southwards dispersal out-of Tanzania, along a latitudinal axis spanning cross Mozambique into South Africa. The resulting gradient of genetic relatedness is accompanied by a gradual reduction in genetic diversity possibly indicative of serial bottlenecks, as well as by a progressive loss of the genetic similarity between East Bantu speakers and Bantu-speaking peoples remaining in West-Central Africa. This increased genetic differentiation, however, cannot be attributed to admixture with resident populations. In fact, the absence of a substantial Khoisan contribution to the genetic make-up of Mozambican Bantu speakers (1-5%) suggests that the migrants had very low levels of admixture with resident populations until they reached the southernmost areas of eastern Africa, where Sotho and Zulu display considerable admixture proportions (24%). Moreover, the dates we obtained for admixture between Bantu speakers and Khoisan groups (∼1165 BP) are remarkably close to the dates for the first archaeological attestations of the presence of Bantu speakers in southeastern Africa. We therefore conclude that our results provide a genetic counterpart to the distribution of Early Iron Age assemblages associated with the Kwale ceramic tradition, which are thought to constitute the material evidence for the southward movement of Bantu speech communities along the Indian Ocean coast.

## Material and Methods

### Population samples

A total of 231 samples from 12 ethnolinguistic groups from Mozambique and three groups from Angola were included in the present study (Fig. 1*A*). Sampling procedures in Mozambique and Angola were described elsewhere (37, 38). All samples were collected with informed consent from healthy adult donors, in collaboration with the Portuguese-Angolan TwinLab established between CIBIO/InBIO and ISCED/Huíla Angola and the Pedagogic and Eduardo Mondlane Universities of Mozambique. Ethical clearances and permissions were granted by CIBIO/InBIO-University of Porto, ISCED, the Provincial Government of Namibe (Angola), and the Mozambican National Committee for Bioethics in Health (CNBS).

### Genotyping and phasing

DNA samples were extracted from buccal swabs and genotyped with the Illumina Infinium Omni2-5Exome-8 v1-3_A1 BeadChip (39, 40), after Whole Genome Amplification (WGA). Of a total of 2,612,357 genomic variants initially typed in 231 samples from Angola and Mozambique, a final set of 200 individuals typed for 1,946,715 autosomal SNPs was retained after applying quality control filters. Haplotypes and missing genotypes were inferred using SHAPEIT2 (41). Geographic locations, linguistic affiliations and sample sizes for all groups are presented in Table S1. Details about DNA extraction, genotyping, haplotyping and quality control filtering are provided in *SI Appendix*.

### Data merging

The newly generated data from Angola and Mozambique were merged with eight publically available datasets (7, 20, 21, 42–46), following the approach described above in *SI Appendix*. The final merged dataset consists of 1,466 individuals from 89 populations typed for 105,286 SNPs (Table S5).

### Genetic data analysis

PCA was performed with the EIGENSOFT v7.2.1 package (24). Unsupervised clustering analysis was done with ADMIXTURE (25) applying a cross-validation (CV) procedure. We performed 20 independent runs for each number of clusters (K) and post-processed and plotted the results with the pong software (47). For PC and ADMIXTURE analyses, SNPs in linkage disequilibrium (r^2^>0.5) were removed using PLINK 1.9 (48), which reduced the newly-generated and merged datasets to 927,435 and 98,570 independent autosomal SNPs, respectively. To assess the relationship between genetic, geographic and linguistic data, we used Procrustes analysis (49), Estimated Effective Migration Surfaces (EEMS) (28) and Mantel tests (50), as detailed in *SI Appendix*. Levels of genetic diversity were assessed by using Haplotype Heterozygosity (HH), Runs of Homozygosity (RoH) and Linkage Disequilibrium (LD), as described in *SI Appendix*. To infer “painting” or copying profiles and quantify the ancestry contributions of different African groups to Bantu-speaking populations of Mozambique, Angola and South Africa, we used CHROMOPAINTER v.2 (32) in combination with the MIXTURE MODEL regression implemented in the GLOBETROTTER software (33). GLOBETROTTER was also used to infer and date admixture events. Details on the application of these methods are provided in *SI Appendix*.

### Linguistic data analysis

We collected published lexical data from 24 languages from Mozambique (10), Angola (3), eastern (9) and southern Africa (2) (Figs. S2 and S10), based on the wordlist published by Grollemund et al. (14) consisting of 100 meanings (Table S3). Using reconstructions provided in the online database Bantu lexical reconstructions 3 (51) in combination with standard methodology from historical-comparative linguistics, we identified 636 cognate sets, and all languages were coded for presence (1) or absence (0) of a particular lexical root. Based on our coded dataset (Table S4), we used the software SplitsTree v4.14.2 (52) to generate a matrix of pairwise linguistic distances (1-the percentage of cognate sharing) and computed Neighbor-Joining networks with 10,000 Bootstrap replicates (Figs. S2*B* and S10).

## Data Availability

The newly generated data will be made available for academic research use through the ArrayExpress database (accession number TBD).

## Acknowledgments

We are grateful to all subjects who participated in this research. Financial support for this work was provided by Foundation for Science and Technology (FCT, Portugal) under the project PTDC/BIA-GEN/29273/2017 and by projects Variabilidade Biológica Humana em Moçambique and STEPS at the Pedagogic University and Eduardo Mondlane University of Mozambique. Genotyping was performed by the SNP&SEQ Technology Platform in Uppsala (www.genotyping.se). The facility is part of the National Genomics Infrastructure supported by the Swedish Research Council for Infrastructures and Science for Life Laboratory, Sweden. The SNP&SEQ Technology Platform is also supported by the Knut and Alice Wallenberg Foundation. The computations were performed at the Swedish National Infrastructure for Computing (SNIC-UPPMAX). AS was supported by the FCT grant SFRH/BD/114424/2016, MGV by POCI-01-0145-FEDER-006821 funded through the Operational Programme for Competitiveness Factors (COMPETE, EU) and UID/BIA/50027/2013 from FCT, AMF by CEECIND/02765/2017 from FCT, and BA by a post-doctoral fellowship of the Fyssen foundation.

