## Supplementary Information for "Mozambican genetic variation provides new insights into the Bantu expansion"

**Supplementary Information Appendix**

### 1. DNA extraction and WGA

DNA was extracted from buccal swabs, using the QIAamp® DNA Micro Kit protocol (QIAGEN®) according to the manufacturer's instructions, and was quantified using the NanoDrop2000 Spectrophotometer (Thermo Fisher Scientific Inc.). Whole Genome Amplification (WGA) was performed with the Illustra GenomiPhi v2 DNA Amplification Kit (General Electric Company-GE Healthcare Life Sciences) according to the manufacturer's instructions. Each batch of WGA was co-amplified with a positive control (GenomiPhi Control DNA) and negative control (distilled water). WGA products were quantified using the Quant-iT™ PicoGreen® dsDNA quantification kit (Invitrogen, P7581) according to the manufacturer's instructions.

### 2. Genotyping, quality control and phasing

In total, 2,612,357 genomic variants from 231 samples were genotyped at the SNP&SEQ Technology Platform at Uppsala University (Sweden), using the Illumina InfiniumOmni2.5Exome-8v1.3 array. The results were analyzed using the Illumina Genome Studio v.2011.1 software and aligned and exported against hg19. Samples had an average call rate of 93.06%. Ten individuals were genotyped by using both genomic DNA and whole genome amplified DNA, showing 99.6% concordance of genotype calls. For quality control (QC) procedures, we used the PLINK v1.9 software (1) and custom scripts to filter out variants and individuals. We excluded 2,233 unmapped variants, 164 insertions and deletions (INDELs), 381 SNPs on the mitochondrial chromosome, 2,218 SNPs on the Y-chromosome, 58,176 SNPs on the X-chromosome, and 2663 on the pseudoautosomal region (PAR). We subsequently removed 65,124 duplicated SNPs, 76,134 A/T or C/G SNPs and 58,489 SNPs with genotype mismatches between genomic and whole genome amplified samples. We additionally excluded 399,977 SNPs with more than 10% missing data and 83 SNPs deviating from Hardy-Weinberg equilibrium in Angola or Mozambique ($p < 0,001$ Fisher's exact test). Overall, the variant filtering yielded 1,946,715 autosomal SNPs. We also excluded seven individuals with >15% missing data and 14 cryptically related individuals, who presented a first-degree or second-degree relationship, as inferred by KING (2).

After these exclusions, and removing the duplicated individuals used to compare genomic and whole genome amplified DNA, the dataset comprised a final number of 200 individuals with an average call rate of 98% (Table S1). Haplotypes and missing genotypes were inferred using SHAPEIT2 (3), with 100 states and 20 MCMC main iterations. To improve the phasing accuracy, the 1000 Genomes Project phase 3 dataset was used as a reference panel and the HapMap phase II b37 reference as genetic map (4).

### 3. Merging with other datasets

The newly generated data from Angola and Mozambique were merged with eight publically available datasets (Table S5) (5–12). Before merging, quality control filtering and missing genotype inference were carried out for each dataset following the approach described above. All SNPs that did not overlap between merged datasets were removed. After filtering and merging, populations with more than 20 sampled individuals were randomly sub-sampled to a sample size of 20. The final merged dataset consists of 1,466 individuals from 89 populations typed for 105,286 SNPs.

#### 4. Genetic diversity

We characterized genetic diversity in Mozambique using haplotype heterozygosity (HH), runs of homozygosity (RoH) and linkage disequilibrium (LD) measured by the squared correlation of allele frequencies ($r^2$) (Figs. 2, S3 and S4). To perform these analyses, we combined the very small sample of Ronga (n=3) with the Tswa (n=6), since the two populations do not display a significant genetic distance ($F_{st}$) and belong to the same linguistic group (Tswa-Ronga). Moreover, to control for uneven sample sizes we downsampled the number of individuals to 5 (the sample size of Mwani, Table S1), before applying a 10% minimum allele frequency cutoff. We repeated this process ten times, calculating each summary statistic in each replicate, and taking the average over replicates as the final estimate.

Expected haplotype heterozygosity was computed using window sizes of 5 SNPs and a step size of 1 SNP, considering each unique 5 SNP haplotype as a separate allele. The mean haplotype heterozygosity for each population was obtained by averaging the heterozygosity across all windows of the genome. These calculations were performed using a custom script with VCFtools (13) and R software (14).

RoHs were calculated with the PLINK v1.9 software (1) under the same parameters previously used by Schelebusch et al. (10). We report both the total number and the total length of RoH, noting that the total length can be more influenced by recent inbreeding than the total number, which probably better reflects long-term population history (15). In any case both statistics are significantly correlated with latitude and with each other (Figs. 2, S3 and S4).

We calculated LD ($r^2$) between pairs of SNPs in sliding windows of 1Mb in each population using PLINK v1.9 (1). To evaluate the LD-decay (Fig. S5), we binned the LD values between pairs of SNPs according to different genomic distance categories (<2 Kb, 2-5 Kb, 5-10 Kb, 10-15 Kb, 15-20 Kb, and 20-25 Kb) and calculated the mean $r^2$ value within each bin.

#### 5. Population structure analyses

#### 5.1. Genotype-based methods

##### 5.1.1 Principal component and unsupervised clustering analyses

Principal component analysis (PCA) was performed with the EIGENSOFT v7.2.1 package (16), removing outliers. Unsupervised clustering analysis was done with ADMIXTURE v1.3.0 (17) applying a cross-validation (CV) procedure and performing 20 independent runs for each number of clusters (K). In Fig. 4*D* and Fig. S6, the results were post-processed and plotted the results with the pong software (18). Maps displaying the distribution of average proportions of ADMIXTURE components (Fig. 4) were plotted with the interpolation plugin of the QGIS v2.18 software, using the Inverse Weighting Distance (IDW) method (19). For PC and ADMIXTURE analyses, SNPs in LD ($r^2$>0.5) were removed with PLINK v1.9 (1) using the option –indep-pairwise 50 10 0.5, which reduced the newly-generated and merged datasets to 927,435 and 98,570 autosomal SNPs, respectively.

#### 5.1.2 Relationship between genetic, geographic and linguistic data

To assess correlations between genetic, geographic and linguistic data we used Procrustes transformation analysis (20), Estimated Effective Migration Surfaces (EEMS) (21) and Mantel tests (22). All sampled individuals belonging to a given ethnolinguistic group were considered to have the same, average geographic coordinates. For computing Procrustes transformation and correlations between PCA results and sampling locations, we used the *procrustes* function implemented in the R software package *MCMCpack* 1.4-4 (23) (Figs. 1, 4, S1 and S8). For effective migration surfaces estimation with the EEMS software, genetic dissimilarities were calculated with the bed2diffs program. The number of demes was set to 200 and we performed six independent runs of 8 million iterations each, discarding the first 4 million iterations as burn-in. The run with highest likelihood was used for a second set of four runs of 4 million iterations each. The *rEEMSplots* R package was used to visualize the data (Fig. 3) (21). For Mantel tests (Table S2), we used pairwise $F_{st}$ between populations as genetic distance (24) calculated with EIGENSOFT v7.2.1 package (16). Geographic distances were calculated as Great Circle Distances using the FIELDS R package v9.0 (25). Linguistic distances were calculated as detailed in Material and Methods - Linguistic data analysis.

### 5.2. Haplotype-based methods

To identify the likely regional sources for the migrations of southwestern and southeastern Bantu-speaking populations, we used CHROMOPAINTER v.2 (26), assuming that the haploid genomes of Angolan, Mozambican and South African Bantu speakers (recipients) were formed by DNA chunks from donors from other populations. To save computation time, we reduced the number of potential donor groups to a subset of 54 populations from the whole merged dataset, consisting of one sample of western European ancestry and 53 groups representing Bantu and non-Bantu speakers from different African regions and cultural traditions. The recipients consisted of 5 Angolan, 12 Mozambican and 2 South African groups, adding up to 19 Bantu-speaking target groups (Table S6). The painting or copy profiles of all individuals from each recipient group were averaged and displayed in a coancestry matrix based on the number of shared DNA chunks between donors and recipients (chunkcounts) (Fig. 5*A*). The differences between the average copy profiles of pairs of recipient populations (X and Y) were quantified using the total variation distance $TVD_{XY}$ (27, 28) (Fig. 5*B*).

To quantify the ancestry contributions of different source populations to the genetic make-up of Angolan, Mozambican and South African Bantu speakers we used the MIXTURE MODEL regression implemented in the GLOBETROTTER software, which uses coancestry matrices generated by CHROMOPAINTER (27–29). In GLOBETROTTER terminology, the recipient populations whose ancestry profiles are being determined are referred to as “targets”, while the source populations used to describe the ancestral composition of targets are referred to as “surrogates”. In addition, GLOBETROTTER requires that copy profiles of target and surrogate populations are first obtained with CHROMOPAINTER from a set of groups referred to as “donor” populations (29). In our analyses, we always used the same set of 73 “donor” groups, consisting of the aforementioned set of 54 populations plus the 19 Angolan+ Mozambican+South African Bantu-speaking groups (Table S6). We then used these populations in different configurations of “surrogate” (source) and “target” (recipient) groups. For example, in Fig. 5*C* we calculated the ancestral profiles of Angolan, Mozambican and South African Bantu-speaking peoples using “target” and “surrogate” groups that correspond to the recipient and source groups displayed in the coancestry matrix of Fig. 5*A*, respectively. In Fig. 6 we attempted to rank the proxies for the ancestry of 2 “target” South African Bantu-speaking groups (Sotho+Zulu) by successively removing from the “surrogate” list populations from areas that had provided best source candidates in a previous run (Table S6).

We also used GLOBETROTTER to identify and date signals of admixture in the history of the studied populations. Table S7 shows the admixture events detected in our Mozambican and Angolan samples, as well as in East African and southern Africans from the pooled datasets. Dates were estimated using a generation time of 29 years (30).

To estimate mutation emission and recombination scaling parameters used in the analyses relying on CHROMOPAINTER, we performed initial runs using 10 iterations of the Expectation-Maximization (EM) algorithm for a subset of five randomly selected chromosomes (1, 6, 11, 16, and 22). The inferred parameters were averaged by chromosome (weighted by their number of SNPs) and then by individuals. These parameters were then used in subsequent CHROMOPAINTER runs on all individuals and chromosomes.

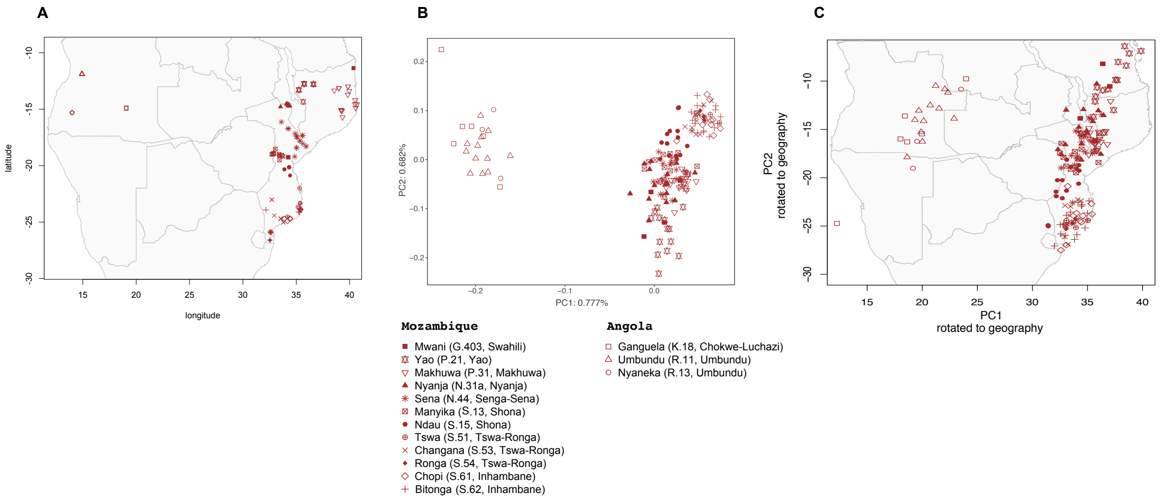


**Fig. S1.** Genetic structure in Angolan and Mozambican populations. (A) Locations of sampled individuals. Geographic subgroups of Bantu languages ("Guthrie zones") following Maho (31) are given in parentheses. (B) Principal components 1 and 2 of Angolan and Mozambican individuals. (C) Principal components 1 and 2 of Angolan and Mozambican individuals rotated to fit geography.

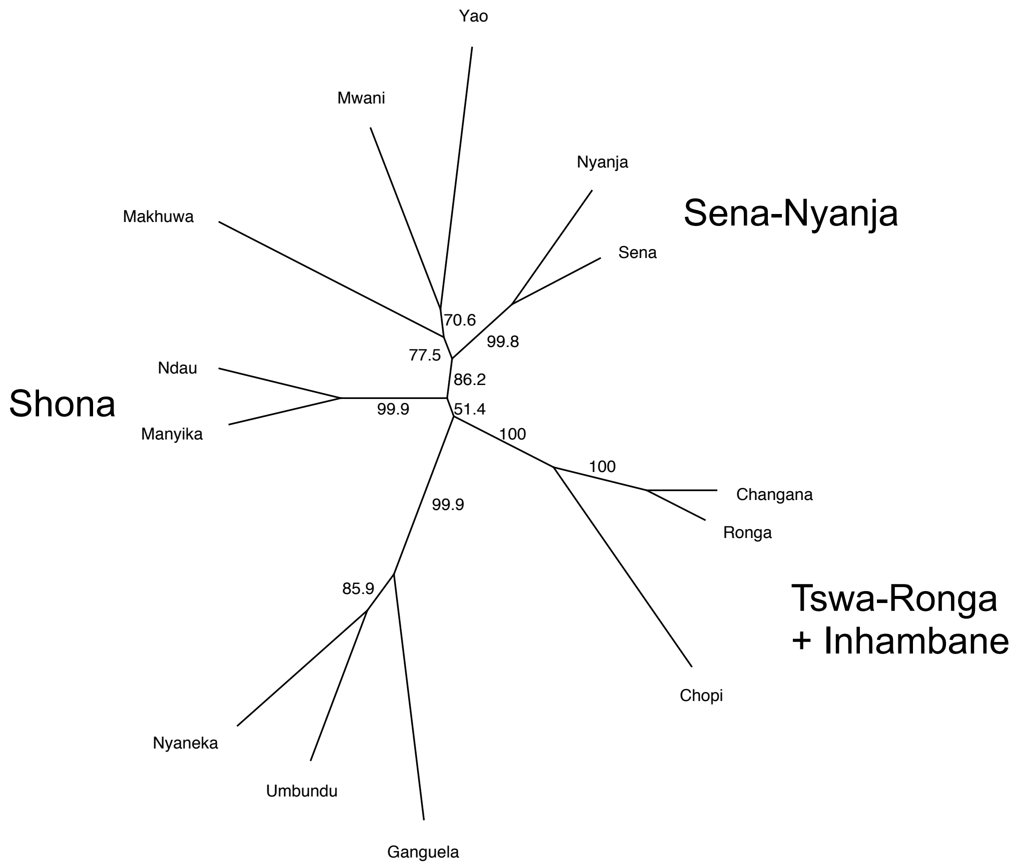


**Fig. S2.** Neighbor-Joining (NJ) network based on linguistic distances among 13 Angolan and Mozambican languages. Values indicate percentages of bootstrap support for internal branches calculated with 10,000 iterations.

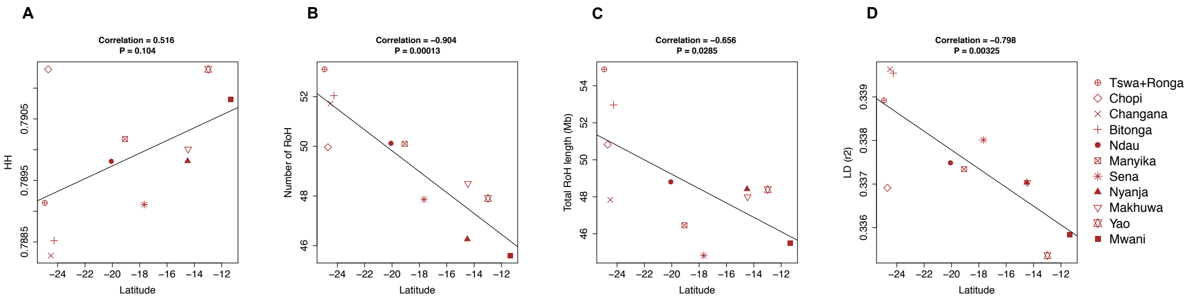


**Fig. S3.** Correlations between different summary statistics of genetic diversity and latitude in Mozambique. (A) Haplotype heterozygosity (HH) (B) Number of runs of homozygosity (RoH) (C) Total RoH length (D) Linkage disequilibrium (LD), as measured by $r^2$.

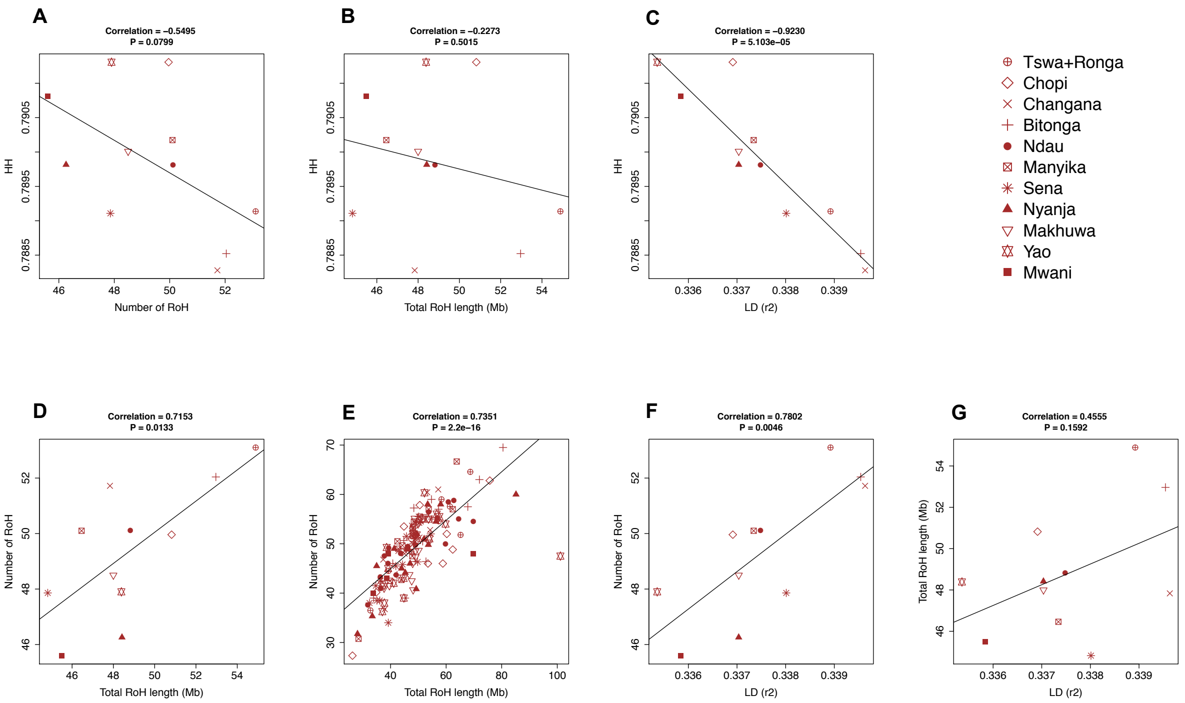


**Fig. S4.** Pairwise correlations between summary statistics of genetic diversity in Mozambique. (A) Correlation between haplotype heterozygosity (HH) and number of runs of homozygosity (RoH). (B) Correlation between HH and total RoH length. (C) Correlation between HH and linkage disequilibrium (LD). (D) Correlation between number of RoH and total RoH length (populations). (E) Correlation between number of RoH and total RoH length (individuals). (F) Correlation between number of RoH and LD. (G) Correlation between total RoH length and LD.

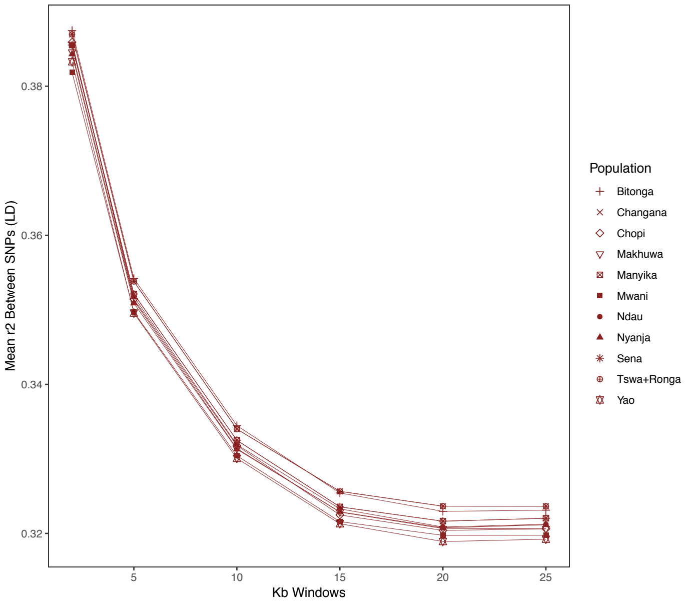


**Fig. S5.** Linkage disequilibrium ($r^2$) decay with physical distance in Mozambican populations.

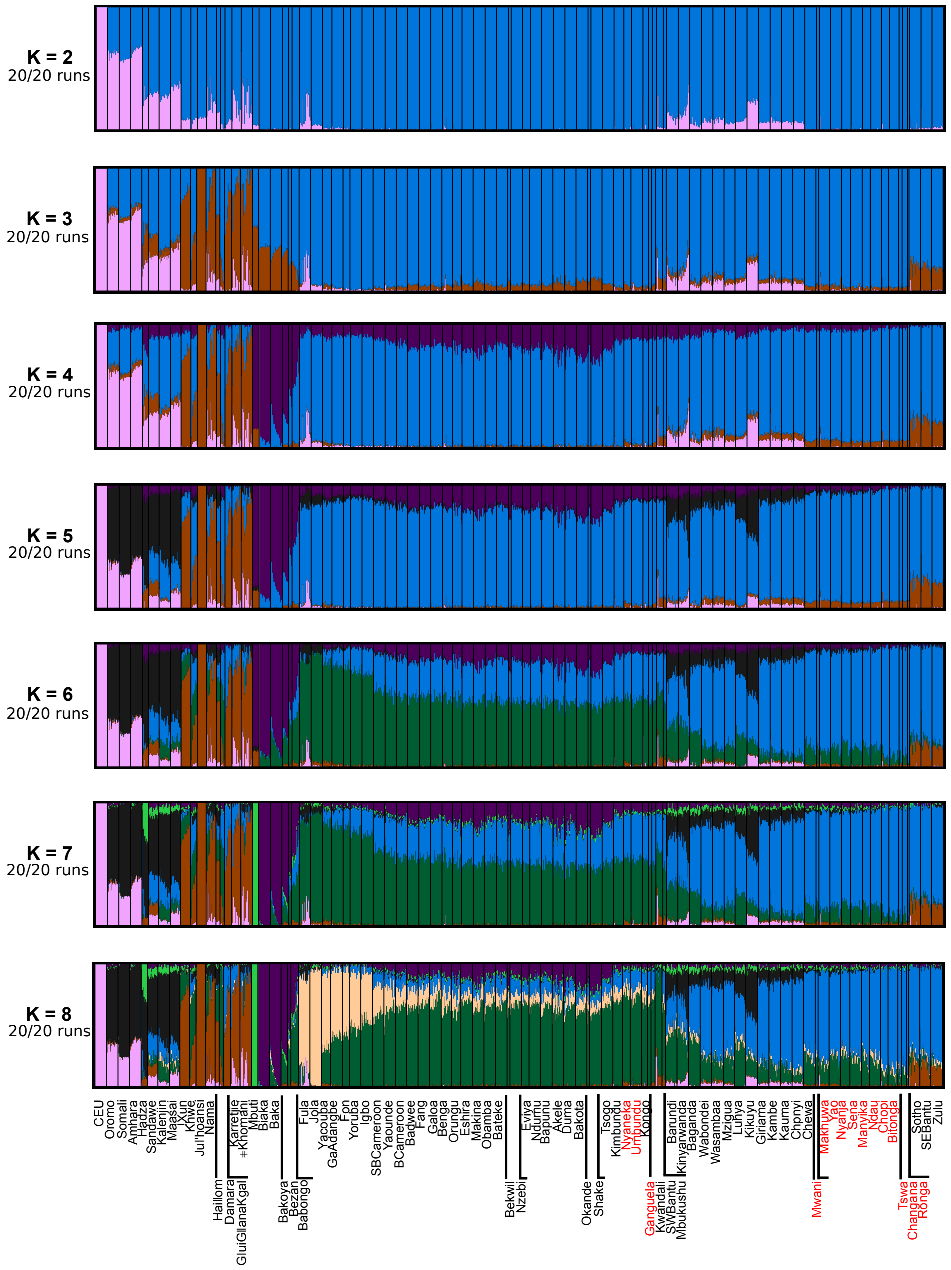


**Fig. S6.** Population structure estimated with ADMIXTURE analysis from K=2 to K=8, with Mozambican and Angolan populations from this study marked in red. Vertical lines represent the estimated proportions of each individual's genotypes that are derived from the assumed K-groups. The lowest cross-validation error was associated with K=6.

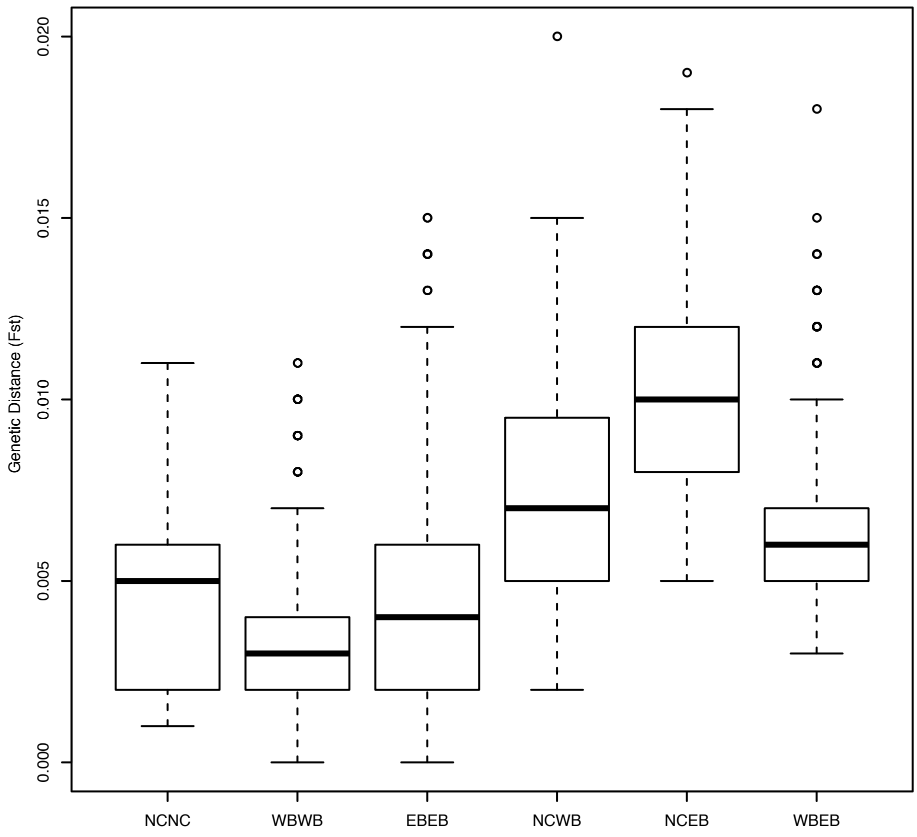


**Fig. S7.** Boxplot graph showing the distribution of genetic ($F_{st}$) distances between different pairwise combinations of populations speaking non-Bantu Niger-Congo (NC), West Bantu (WB) and East Bantu (EB) languages.

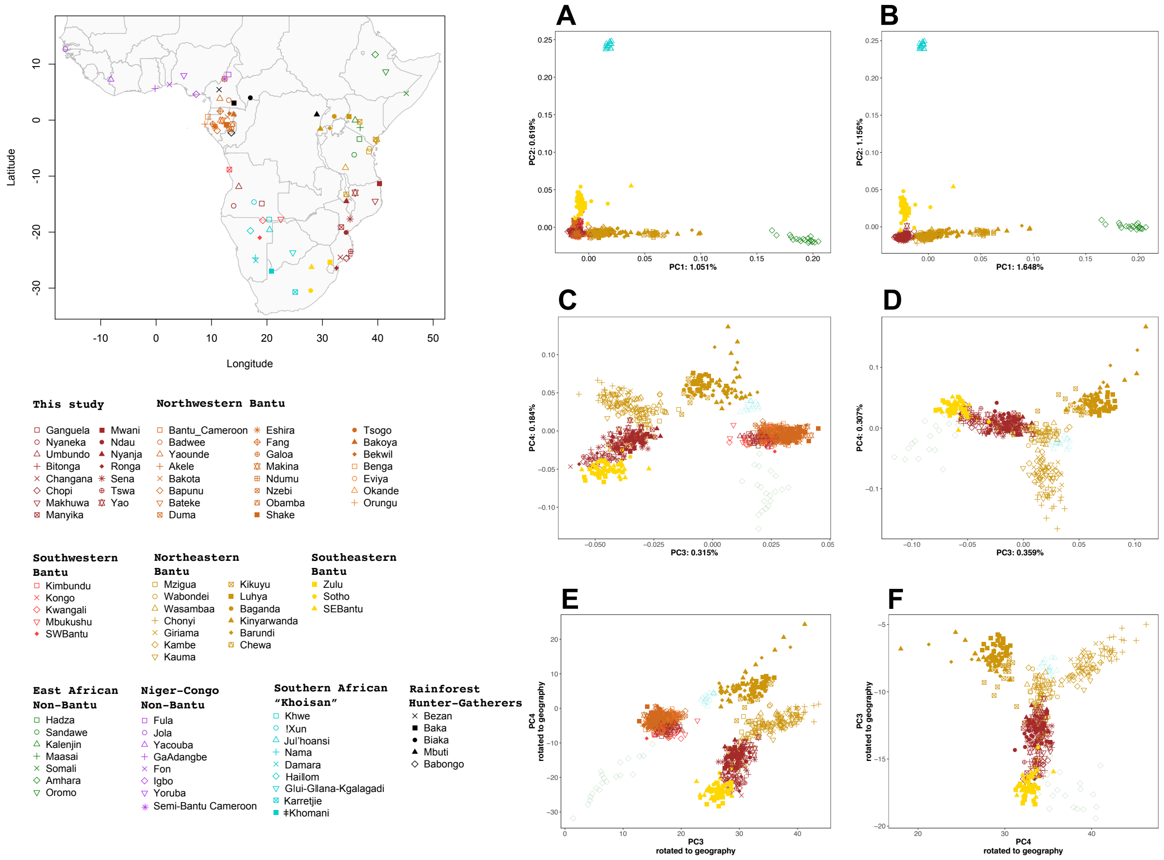


**Fig. S8.** PC analyses of all Bantu (A, C, E), and East Bantu speakers (B, D, F), together with one representative group of southern Africa "Khoisan" (Jul'hoansi) and one representative group of non-Bantu eastern Africans (Amhara). Plots E and F were rotated to fit geography using Procrustes analysis (E: Procrustes correlation: 0.76; P< 0.001; F: Procrustes correlation: 0.44; P<0.001).

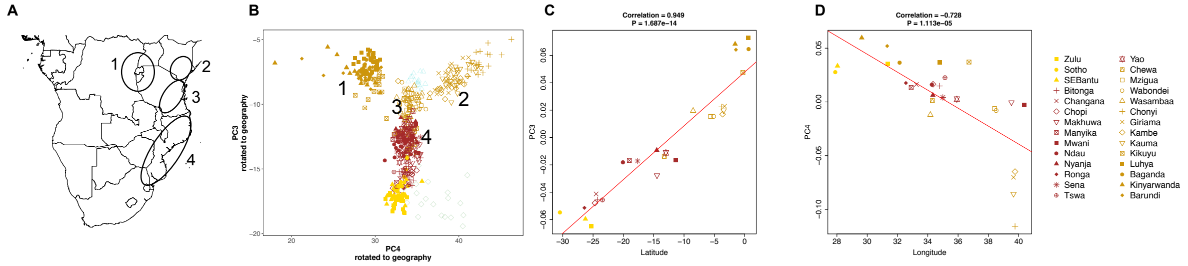


**Fig. S9.** Genetic structure in East Bantu-speaking populations. (A) Geographic location of 4 groups of East Bantu-speaking populations. (B) PC3vsPC4 plot of East Bantu speakers, together with one representative group of southern Africa “Khoisan” (Ju|’hoansi) and one representative group of non-Bantu eastern Africans (Amhara). (C) Correlation between average PC3 scores of East Bantu speakers and latitude. (D) Correlation between average PC4 scores of East Bantu speakers and longitude. Note that Ju|’hoansi and Amhara were excluded from the correlation plots.

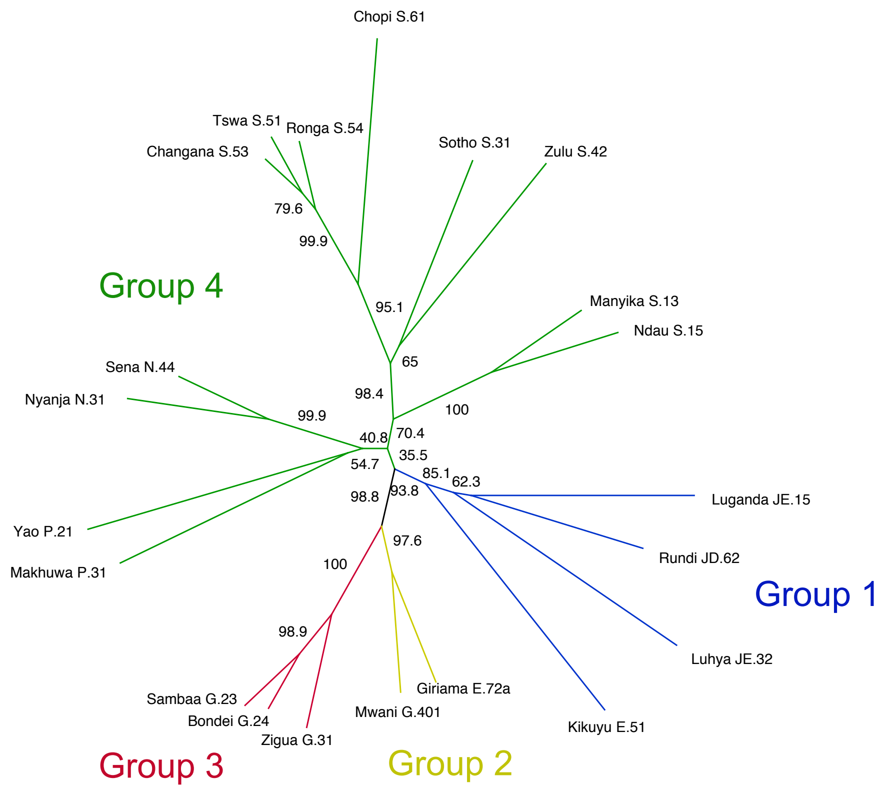


**Fig. S10.** NeighborJoining (NJ) network of linguistic distances among 21 East Bantu languages. Values indicate percentages of bootstrap support for internal branches calculated with 10,000 iterations.
